# A workflow for absolute apoplastic pH assessment during live cell imaging in plant roots

**DOI:** 10.1101/2025.06.06.658240

**Authors:** Ann-Kathrin Rößling, Niklas Mayle, Elke Barbez

## Abstract

Apoplastic pH is a key regulator of plant development and environmental responses, influencing processes such as cell expansion, nutrient uptake, and intercellular signaling. Accurate tools for measuring absolute pH values at high spatial resolution are therefore essential, yet limiting. Here, we present a novel calibration-based workflow for the *in vivo* quantification of absolute apoplastic pH using the fluorescent pH indicator HPTS. While HPTS has previously been used primarily to track relative pH changes, our novel methodology enables precise absolute pH measurement through a simplified calibration strategy and a tailor-made Fiji Plugin. This approach offers a non-invasive, reproducible tool for investigating absolute extracellular pH with high spatial resolution, expanding the methodological toolbox available to plant physiologists.

## INTRODUCTION

The pH of the plant apoplast plays a central role in regulating growth, development, and environmental adaptation (Du et al., 2020; Gámez-Arjona et al., 2022; Geilfus, 2017). Classical models, such as the acid growth theory, highlight how apoplastic acidification in Maize coleoptiles promotes cell elongation by activating wall-loosening enzymes like expansins (Rayle & Cleland, 1970; Hager, 2003). In addition, auxin-triggered apoplast alkalinization steers root bending during gravitropism in Arabidopsis (Barbez et al., 2017). Abiotic stresses, including salinity and drought, also induce changes in apoplastic pH that help initiate systemic defense responses (Geilfus, 2017).

Despite its physiological importance, accurately quantifying apoplastic pH in living tissues remains technically challenging. In previous work, we demonstrated that 8-hydroxy-pyrene-1,3,6-trisulfonic acid trisodium salt (HPTS) is a reliable and effective dye for the relative assessment of apoplastic pH in *Arabidopsis thaliana* roots (Barbez et al., 2017). HPTS is a cost-effective, non-toxic fluorescent indicator that passively diffuses into the root apoplast and performs well in live-cell imaging applications. Like other pH-indicators, HPTS exists in a dynamic balance between its protonated and deprotonated states. Interestingly, the protonated form has an excitation wavelength peak at 405 nm, whereas the deprotonated form absorbs maximally at 465 nm. Regardless of the protonation state, both forms emit fluorescence at around 514 nm. By exciting the sample at both 405 nm and 465 nm separately and measuring the resulting fluorescence intensities, it becomes possible to estimate the relative abundance of each form. The ratio of these two signals provides a reliable readout that correlates with the surrounding pH. So far, HPTS enabled many researchers to assess relative pH in the apoplast of Arabidopsis roots (Dünser et al., 2017; Wang et al., 2022; Dang et al., 2022, Liu et al., 2022; Li et al., 2021; Lin et al., 2021; Rößling et al., 2024; Wang et al., 2025).

The assessment of relative pH dynamics has dissected substantial plant physiological processes in the past and will continue to do so in the future. However, the study of specialized biophysical and biochemical processes may benefit from the measurement of absolute apoplastic pH values. The activity of enzymes, the behavior of pH-sensitive binding sites, and the charge state of biomolecules all depend on defined pH thresholds. Subtle differences in proton concentration can influence molecular conformation, reaction rates, and interaction affinities. Spatially resolved measurements of absolute pH are therefore essential for linking extracellular pH dynamics to their functional consequences in plant physiology. Using HPTS, absolute pH approximations are currently possible but require the generation of an extensive *in vivo* calibration curve in each experiment to account for variability (Barbez et al., 2017). Generating an *in vivo* calibration curve is therefore helpful, but laborious and time-consuming.

Here, we introduce a novel calibration strategy that enables the quantification of absolute apoplastic pH values *in vivo*. This approach relies on a normalization step that reduces experiment-to-experiment variability. The simplified workflow removes the need for generating multiple *in vivo* pH standards while maintaining measurement accuracy. To support image analysis, we provide a user-friendly Fiji plugin that transforms HPTS ratio images into absolute pH maps, where each pixel reflects the local extracellular pH.

## RESULTS

### Absolute pH assessment in vitro

To characterize the pH-dependent behavior of HPTS under controlled conditions, we prepared liquid ½ MS medium supplemented with 1 mM HPTS and adjusted the pH across a range from 5.0 to 7.0. Fluorescence emission at 514 nm was recorded upon excitation at 405 nm (protonated form) and 465 nm (deprotonated form) using a microplate spectrophotometer. Signal intensities were measured across different detector gain settings to assess the robustness of the ratiometric signal across variable imaging conditions. For each sample, we calculated the 465/405 fluorescence intensity ratio and implemented a min-max normalization strategy to account for variation in signal strength due to detection gain, laser power, or sample handling.

Normalization was performed according to the formula:

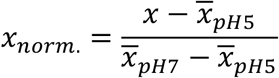

*x*= measured 465 nm/405 nm fluorescence intensity ratio for any sample

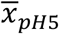= mean ratio at pH 5.0 (lower calibration point)

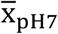= mean ratio at pH 7.0 (upper calibration point)

X_norm_= normalized fluorescence ratio between 0 and 1

This normalization spreads all values between 0 and 1, allowing comparability across independent measurements. Plotting the normalized HPTS ratios against the known pH of each solution revealed a strong, non-linear correlation (Fig. 1a). Regression analysis yielded a third-order polynomial that accounted for 99.76% of the variance, suggesting an excellent fit to the data (Fig. 1a).

**Figure 1:**
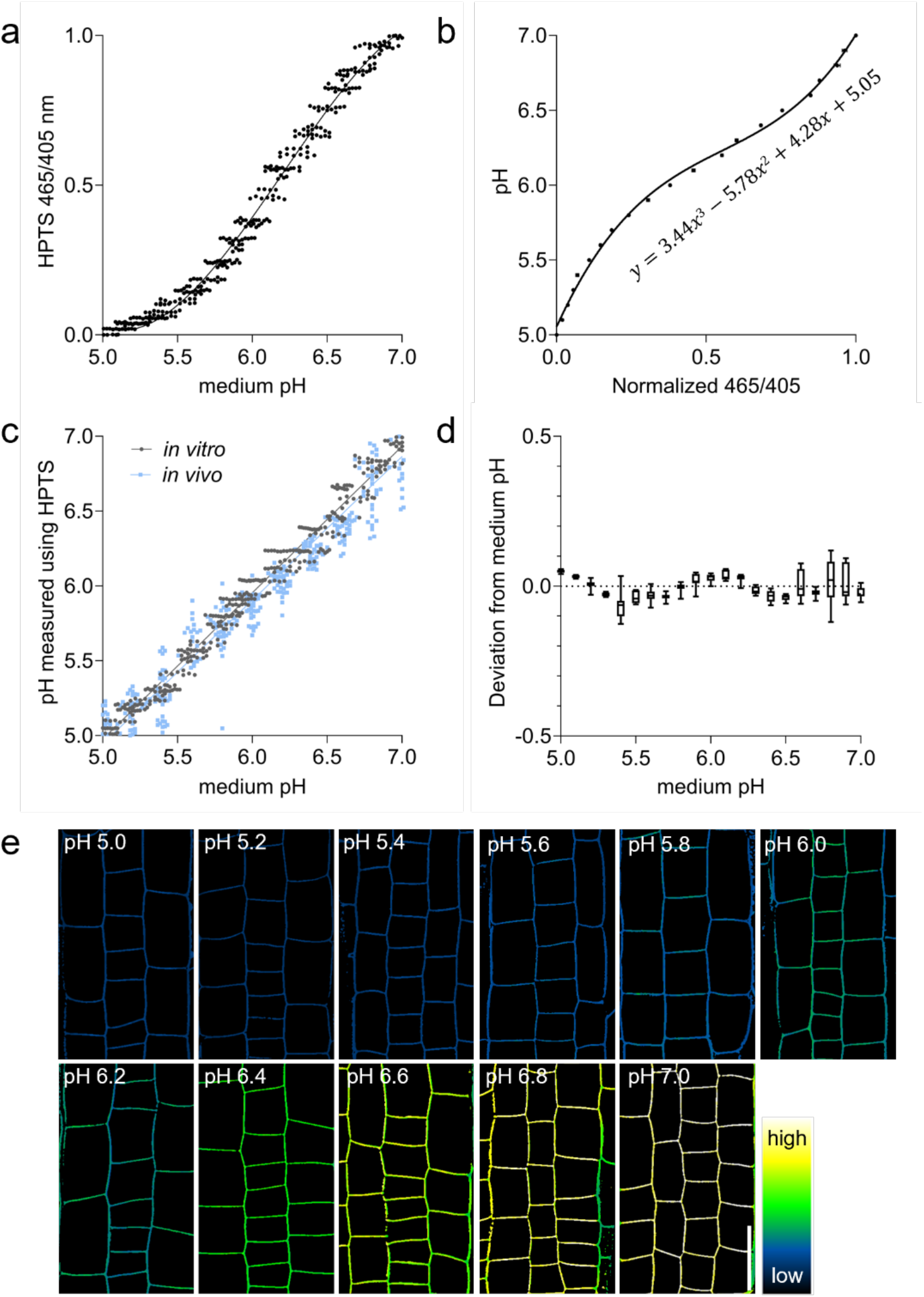
HPTS calibration *in vitro* and *in vivo*. (a and b) *In vitro* HPTS characterization. (a) Graph depicts the min-max normalized 465/405 fluorescence ratio values of HPTS-supplemented liquid medium with a pH across a range from 5.0 to 7.0 as measured using a microplate spectrophotometer. The regression curve with R^2^= 0.9976 is shown. (b) Graph shows the medium pH plotted against the average normalized 465/405 values from (a). The regression equation that converts normalized HPTS ratios into absolute pH values is shown. (c) The pH measured *in vitro* and *in vivo* via HPTS normalization and conversion is plotted against the medium pH. (d) Box-plots depict the deviation of HPTS measured pH *in vitro* from the original medium pH. (e) *In vivo* HPTS characterization. Representative confocal images of seedlings incubated for 10 min and subsequently imaged in HPTS supplemented medium with varying pH between pH5 and pH7. Scale bar = 25 µm, n = 9 – 12.

Since the normalized HPTS ratios robustly correlate with pH, this data set enabled us to formulate an equation that converts normalized HPTS ratios to absolute pH values (Fig. 1b).

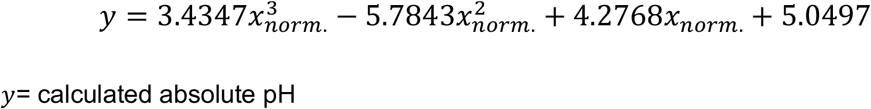

Using this equation, we calculated pH values from normalized ratios and confirmed that the predicted values closely matched the actual pH of the samples (Fig. 1c).

We next calculated how much the calculated pH values deviate from the original medium pH. This analysis helps us understand how well HPTS can be used to differentiate between small changes in pH levels (Fig. 1d). The absolute deviation from the original medium pH was on average 0.034 +/-0.001 pH units. We do observe slightly higher deviations above pH 6.5, presumably due to the reduced buffer capacity of MES at higher pH. Overall, this data indicates that HPTS is able to reliably distinguish pH differences smaller then 0.1 pH units (Fig. 1d).

### Absolute pH assessment in vivo

To test our calibration strategy in a biological context, we incubated *Arabidopsis thaliana* Col-0 seedlings for 10 min in HPTS-supplemented liquid medium adjusted to pH values between 5.0 and 7.0. Via confocal microscopy, we imaged the protonated (λ_em_= 405 nm) and deprotonated (λ_em_= 465 nm) HPTS versions in 2 separate channels.

The resulting fluorescence images were processed using our custom-made ratiometric Fiji plugin (Barbez et al., 2017; Rößling et al., under revision), which provides an output image where each pixel depicts the ratio of the 465nm/405nm signal intensities (Fig. 1e). In order to normalize the HPTS intensity ratios in analogy to the *in* v*itro* approach above, we measured the average HPTS intensity ratios of the roots incubated in pH5 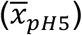 and pH7 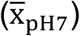.We subsequently measured the average HPTS intensity ratios of the roots incubated in all pH solutions and normalized those values using the same min-max approach as *in vitro.* The resulting normalized ratios were then used to calculate absolute pH values via the calibration equation derived from the *in vitro* dataset.

Plotting the calculated apoplastic pH values against the pH of the incubation medium revealed a strong correspondence between predicted and actual values (Fig. 1c). This confirms that the normalization and calibration approach developed *in vitro* is applicable to *in vivo* imaging and can reliably estimate extracellular pH in living plant tissues.

### The Fiji Plugin

To facilitate data analysis, we developed a custom Fiji plugin that converts ratiometric HPTS images into absolute apoplastic pH maps (Fig. 2). The plugin requires a pre-processed 32-bit grayscale ratio image as input (Fig. 2a), in which each pixel represents the fluorescence intensity ratio between HPTS excited at 465 nm and 405 nm (465/405) (Rößling et al., under revision; Barbez et al., 2017). Users can choose to normalize the image using either manually entered reference values or by loading calibration images from samples imaged at pH 5.0 and pH 7.0 (Fig. 2b). The plugin performs min-max normalization on every pixel of the ratio image using the following equation:

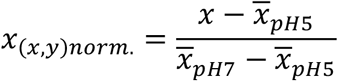

**Figure 2.**
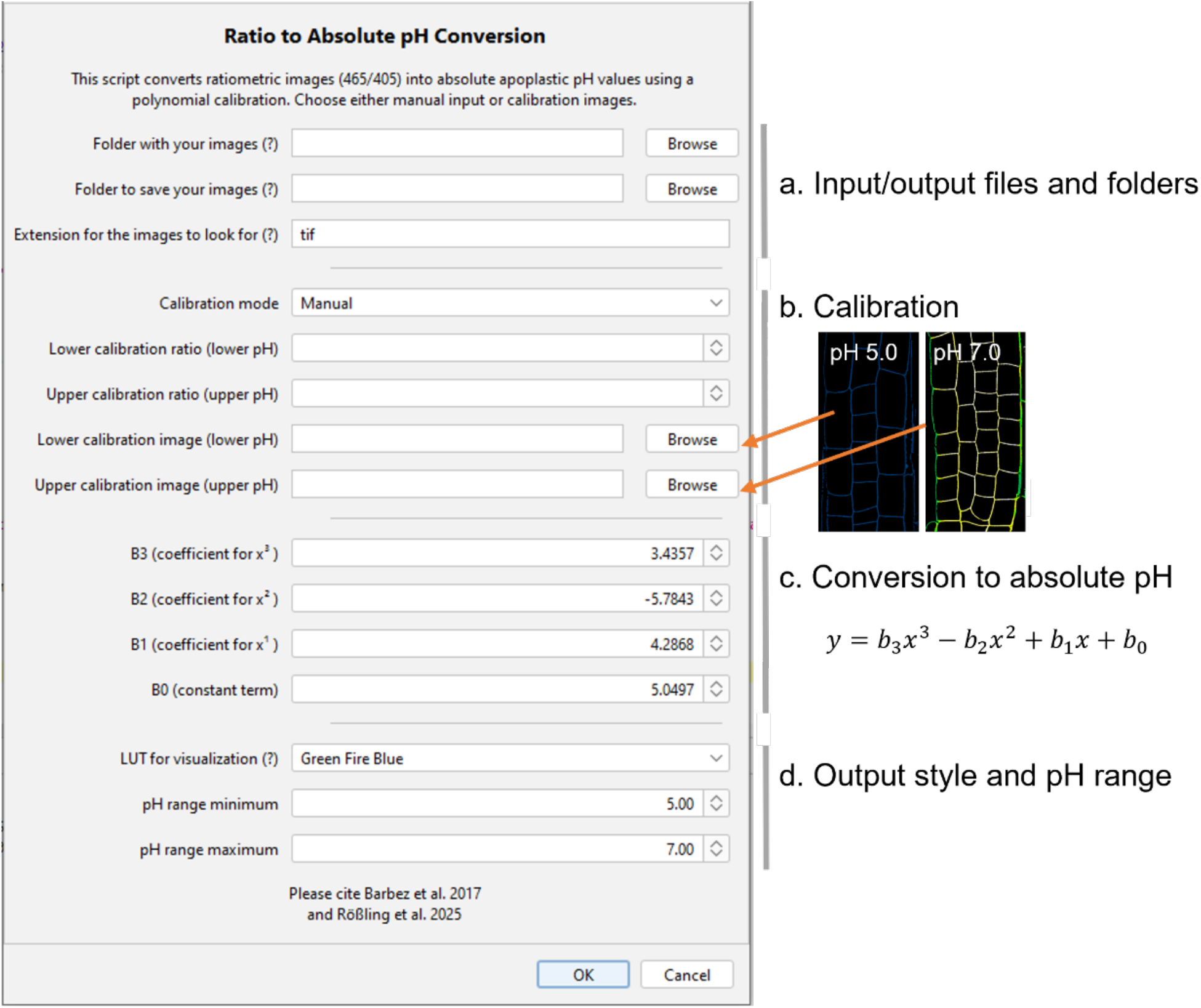
Converting HPTS ratios into absolute pH values: A Fiji Plugin. Our custom-made Fiji macro enables users to convert HPTS ratio images into images depicting absolute pH values. (a) Define the location of the input files (HPTS ratio files) and the location where the output images should be stored. Define the file extension of the input files. (b) Decide to enter calibration values manually (manual) or to upload representative calibration images (from calibration images). The macro can consider only one option. (c) Enter the polynomial equation coefficients from our default analysis (B3 = 3.4347; B2 = -5.7843; B1 = 4.2768; B0 = 5.0497) OR from an adjusted *in vitro* HPTS calibration and regression analysis. (d) Choose a color code (LUT) that will depict absolute pH values, define the range of pH values that will be shown.

(*x, y*)= pixel in position x, y

*x*_(*x, y*)_= 465 nm/405 nm intensity ratio per pixel

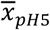= mean 465 nm/405 nm intensity ratio of the loaded calibration image OR manual value

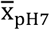= mean 465 nm/405 nm intensity ratio of the loaded calibration image OR manual value

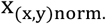= normalized fluorescence ratio per pixel between 0 and 1

Subsequently, it converts each pixel’s normalized value into an absolute pH using the calibration function derived from *in vitro* experiments (Fig. 2c).:

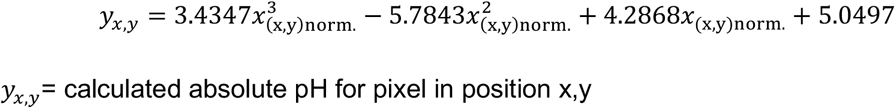

For users working with different buffer systems or experimental conditions, the plugin offers the option to input a custom calibration function. By performing their own *in vitro* calibration using HPTS in the relevant buffer system, users can determine a polynomial regression that best fits their data. The plugin interface allows manual entry of the corresponding coefficients (*b_0_* to *b_3_*), enabling flexible adaptation to alternative calibration curves and ensuring accurate pH conversion across diverse experimental setups (Fig. 2c).

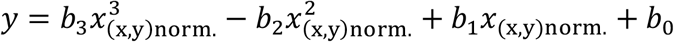

The plugin outputs a 32-bit image in which absolute pH values are color-coded using a user-defined lookup table (LUT) applied across a specified pH range (Fig. 2d). These images support precise quantification of apoplastic pH and can be further analyzed in Fiji, allowing users to extract pH values from defined regions of interest.

### Workflow for absolute apoplastic pH assessment

In order to assess approximate absolute apoplastic pH values we created another Fiji plugin. In order to use this Fiji Plugin, users have to image two calibration samples per experiment: one incubated in HPTS-supplemented medium at pH 5.0 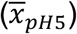 and the other at pH 7.0 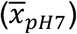,each imaged using identical confocal settings. These reference roots — typically 3 to 5 wild-type seedlings per pH — serve to define the 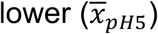 and upper 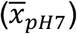 bounds of the 465/405 ratio range for normalization. Since HPTS signal intensities are sensitive to imaging depth, laser power, and detector gain, it is important to define calibration values for each experiment and each tissue type of interest. This approach allows the normalization step to correct for experimental variation, enabling reliable conversion of ratiometric HPTS data into absolute apoplastic pH values (Fig. 3).

**Figure 3:**
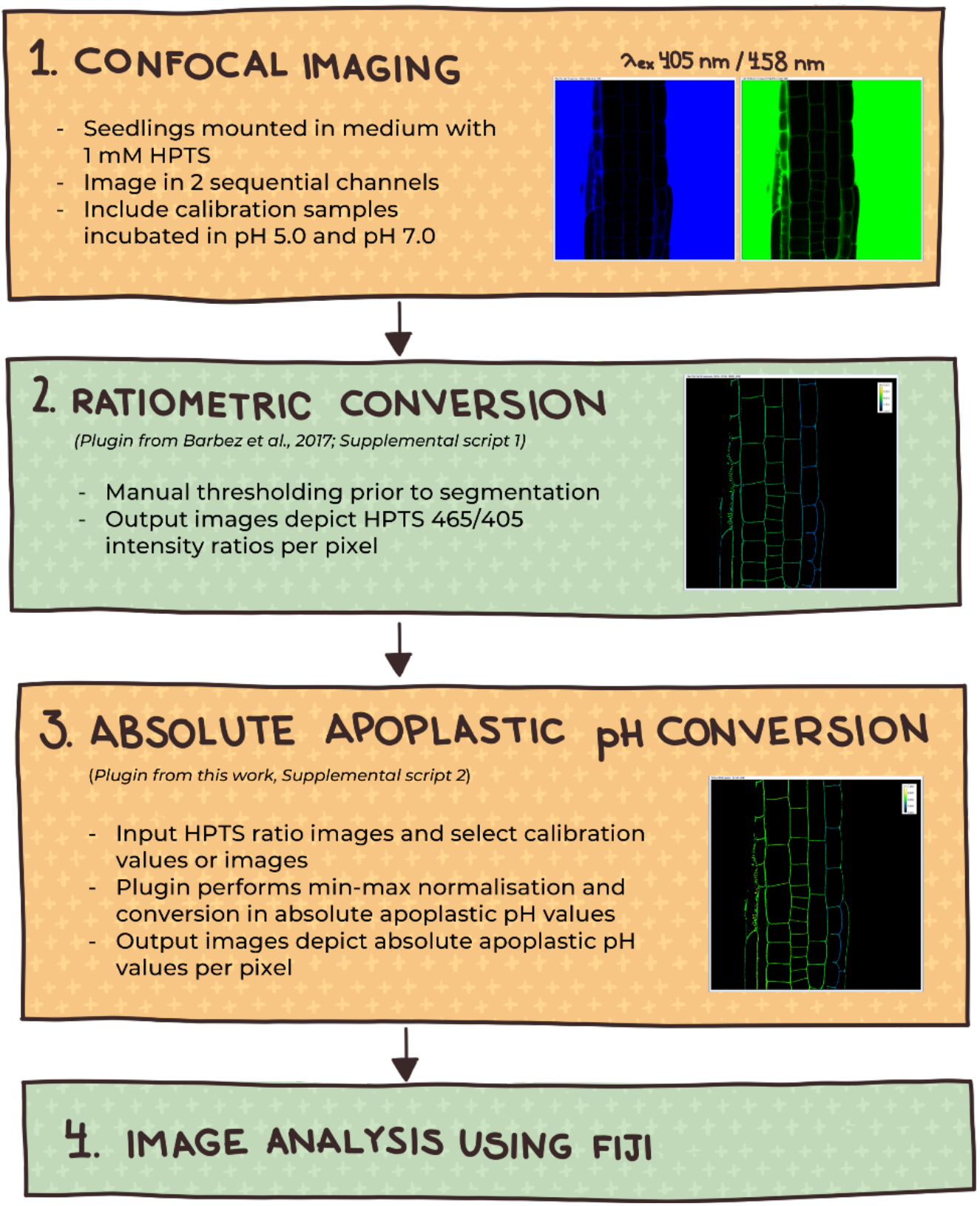
Workflow of absolute apoplastic pH assessment using HTPS. The first step **(1)** is confocal imaging using 1 mM HPTS for mounting and staining. The imaging setup should allow for imaging sequentially in two channels to get the raw images. **(2)** Using the Ratiometric Conversion Fiji Plugin the raw images can be transformed into a 32-bit ratio image. These images can be used for determining pH differences relative to each treatment or line. For absolute pH values, the Fiji plugin for absolute pH conversion is needed **(3)**. This plugin converts the beforehand created ratio images and applies our *in vitro* calibration equation as well as implementing the calibration points pH 5.0 and pH 7.0 to correct for experimental variation. The process results in 32-bit images from which image analysis and quantification in tissue of interest can be performed in Fiji **(4)**.

### Dynamic regulation of absolute apoplastic pH in roots

To verify the functionality of our calibration-based workflow, we tested the effect of several pharmacological treatments on the absolute apoplastic pH in the root epidermis of six-day-old Arabidopsis seedlings. To calibrate our images, we incubated Col-0 wild-type seedlings for 10 min in HPTS-supplemented liquid ½ MS medium of pH 5.0 and pH 7.0. The obtained images were ratiometrically converted using our previously described Fiji ratiometric conversion script (Rößling et al., under revision; Barbez et al., 2017) as well as the Fiji Plugin for absolute pH annotation.

Fusicoccin activates, while DCCD inhibits proton pump activity at the plasma membrane, resulting in a lower and higher apoplastic pH, respectively (Marré, 1979). Fusicoccin resulted in a 0.6 pH unit decrease, while DCCD increased the apoplastic pH with 0.25 pH units compared to mock-treated seedlings. In Barbez et al., we showed that the phytohormone auxin triggers a fast apoplast alkalization (Barbez et al., 2017). Here, we are able to show that IAA treatment results in a 0.2 pH unit increase in apoplastic pH (Fig. 4).

**Figure 4:**
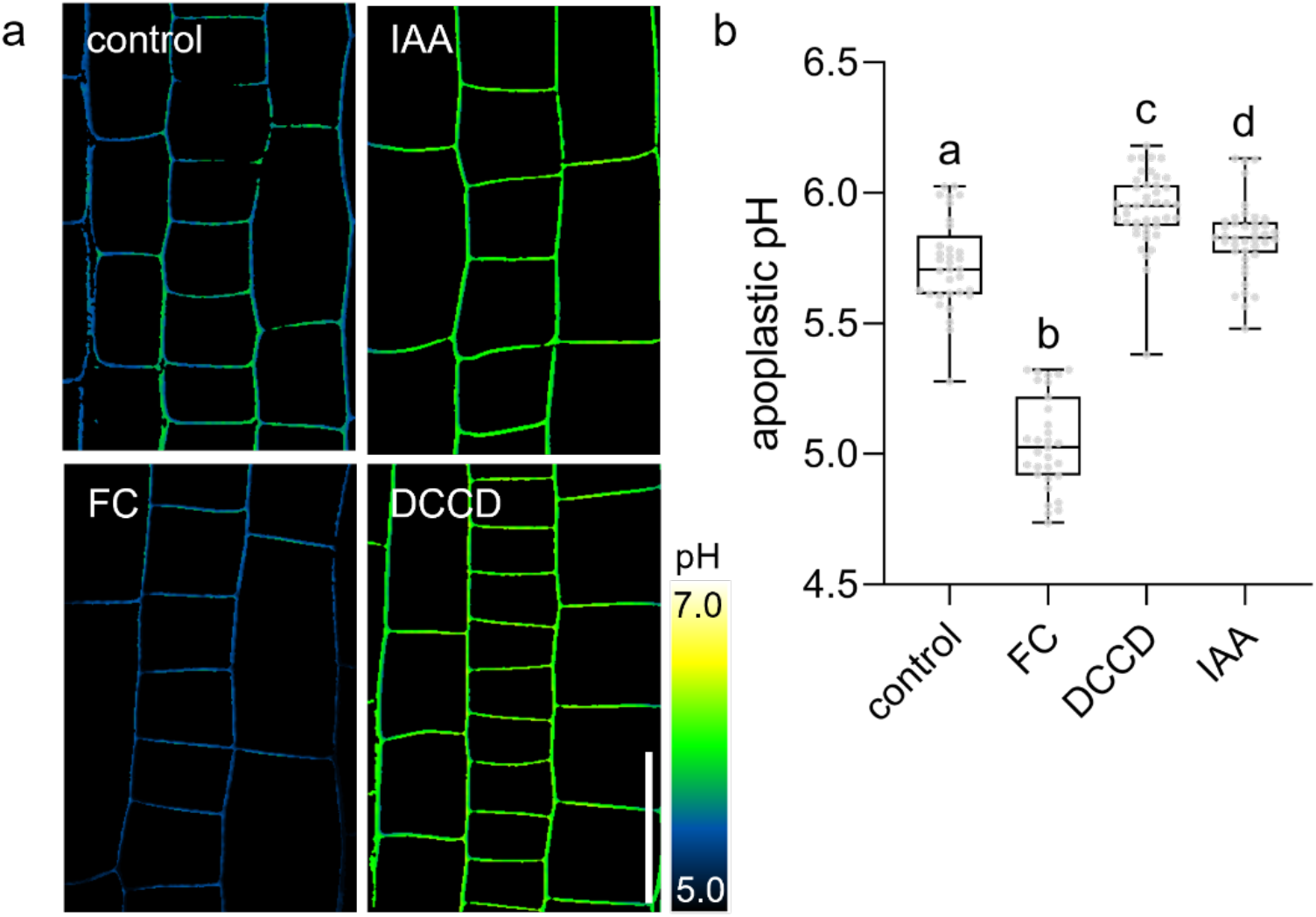
Dynamic regulation of absolute apoplastic pH in Arabidopsis roots. (a) HPTS-based assessment of apoplastic pH in roots 6-day-old of Col-0 wild-type seedlings, treated with 5 µM FC for 3 h, 5 µM DCCD for 3 h or 250 nM IAA for 1 h or solvent control. n = 8 – 12 roots per line/condition. (b) Boxplots depict absolute apoplastic pH from treatments shown in (a). Scale bar = 25 µm. Boxplots: Box limits represent the 25^th^ and 75^th^ percentile, and the horizontal line represents the median. Whiskers display min. to max. values. Representative experiments are shown. Statistical significance was determined using a one-way ANOVA with a Tukey Post Hoc multiple comparisons test (p<0.05, letters indicate significance categories).

## CONCLUSION AND OUTLOOK

Apoplastic pH is a highly dynamic parameter that steers a plethora of plant developmental aspects. The assessment of relative pH dynamics in the apoplast has been very valuable for plant biological research so far. However, biochemical and biophysical applications often require information on absolute apoplastic pH values at a cellular resolution.

Our workflow addresses this gap by combining a simple yet robust normalization strategy with a flexible Fiji plugin, making absolute apoplastic pH imaging more accessible and reproducible. The modular design of the Fiji plugin allows users to tailor the calibration function to their own imaging conditions or buffer systems, extending the method’s flexibility. The ability to derive absolute pH maps at cellular resolution provides a bridge between visual observations and biochemical interpretation. Since HPTS is not toxic and bleach resistant, our workflow can be used to measure apoplastic pH assessment during root growth over several hours or days. As plant biologists increasingly recognize the apoplast as an active regulatory compartment, accessible tools like this will be key to decoding its complex contributions to adaptive plant development.

## MATERIAL AND METHODS

### Plant growth conditions

All experiments were carried out in *Arabidopsis thaliana*, ecotype Col-0. After surface sterilization using 70% and 100% ethanol, the seeds were sown on Murashige and Skoog (MS) plates with 1% sucrose and a pH of 5.7, stratified for 2 days at 4°C in darkness. Afterwards, they were grown vertically in a long-day regime (16 h light, 8 h darkness) at 21°C/18°C.

### Chemicals

Chemicals were dissolved in dimethyl sulfoxide (DMSO), which then also served as a solvent control. Fusicoccin (FC) (Sigma, MO, USA), from *Fusicoccum amygdali,* was used as a proton pump activator, whereas we used N,N’-Dicyclohexylcarbodiimide (DCCD) (Sigma (MO, USA) to chemically inhibit proton pumps. For auxin treatment, 3-Indoleacetic acid (IAA) was utilized (OlChemIm, Czech Republic).

### In vitro characterization of HPTS fluorescence

To assess the spectral behavior of HPTS in response to pH, we performed an *in vitro* calibration using HPTS in liquid plant growth medium. Liquid ½ Murashige and Skoog (MS) medium was prepared and adjusted to pH values ranging from 5.0 to 7.0 in 0.1 pH unit increments. All solutions were autoclaved and cooled to room temperature prior to use. HPTS (8-hydroxy-pyrene-1,3,6-trisulfonic acid trisodium salt; Sigma-Aldrich) was added to each solution to a final concentration of 1 mM. For each pH condition, 200 µL of HPTS-supplemented medium was transferred in quadruplicate into a black, flat-bottom 96-well microplate. Fluorescence intensity was measured using a microplate reader with excitation wavelengths at 405 nm and 465 nm, and emission recorded at 514+/-8 nm for both channels. The ratio of fluorescence intensities (465/405) was calculated for each replicate. Ratio values were min-max normalized using the mean ratios from the pH 5.0 and pH 7.0 samples, respectively. The normalized ratio (x_,norm_) was then plotted against the known pH values, and a third-order polynomial regression was applied to generate a calibration curve. The quality of the fit was assessed using the coefficient of determination (R^2^), with values above 0.95 considered acceptable. A second regression was performed by plotting pH as a function of x_,norm_to derive an equation for the conversion of normalized fluorescence ratios into absolute pH values.

### Confocal microscopy

For imaging, an upright Leica TCS SP8 FALCON FLIM confocal laser scanning microscope equipped with a Leica HC PL APO Corr 63x 1.20 water immersion CS2 objective was used. Apoplastic pH assessment using 8-hydroyypyrene-1,3,6-trisulfonic acid trisodium salt (HPTS; Sigma-Aldrich) was carried out as introduced by Barbez et al., 2017 and updated by Rößling et al., under revision. The protonated form of HTPS was excited at 405 nm using a diode UV laser at 0.5% intensity, whereas the deprotonated form was excited at 458 nm utilizing an Argon laser, running at 30% intensity. Images were obtained in sequential scanning mode. As detectors, PMTs were used. The PMT gain stayed constant during imaging.

For *in vivo* imaging of apoplastic pH in *Arabidopsis thaliana* roots for the calibration curve, six-day-old seedlings were incubated in HPTS-supplemented ½ Murashige and Skoog (MS) liquid medium with different pH values between 5.0 – 7.0 in steps of 0.2 prior to confocal imaging. The treatments were carried out as described in the figure legend. Confocal images were acquired sequentially using excitation at 405 nm and 465 nm and a common emission window at 514 nm.

For image analysis, the previously developed Fiji plugin for image conversion (Barbez et al., 2017, Supplemental Script 1) and the here presented plugin for absolute pH conversion (Supplemental Script 2 https://doi.org/10.5281/zenodo.15599806) were used. The obtained pictures depicting absolute apoplastic pH values were further quantified using Fiji.

### The Fiji plugin

The here developed “Ratio to Absolute pH Conversion” plugin is available at Zenodo: https://doi.org/10.5281/zenodo.15599806. To convert the 32-bit ratiometric HPTS images into absolute apoplastic pH value images, we used the here developed Fiji plugin implemented in Jython (Python for ImageJ). The macro prompts the user to input lower and upper calibration values (i.e., mean fluorescence intensity ratios at pH 5.0 and pH 7.0, respectively), or select representative pictures of seedlings incubated in liquid medium of pH 5.0 and pH 7.0.

The calibration values (manual mode) or the mean intensities of the calibration images (automated mode) will be used for normalization. For the automated mode, the plugin retrieves the mean grey value of the uploaded calibration images of the upper and lower calibration points. The mean gray value is obtained by calculating the sum of all pixels and dividing it by the number of pixels using Fiji’s image statistics. For a very specific region of interest we recommend the manual mode for calibration. The values are used for the min-max normalization of the ratiometric data, distributing them between 0 and 1. The normalized values are then converted to absolute pH using a third-order polynomial regression equation derived from *in vitro* HPTS characterization. If a custom calibration equation, for instance, for a different buffering system, is desired, the coefficients can be changed. Users can then choose a LUT, which the calibration bar will display with the absolute pH range. It has user-defined limits, with the default being pH 5.0 and 7.0.

The resulting images display absolute pH values per pixel and are saved in 32-bit format for further analysis. For displaying the images, they need to be converted to RGB images first.

## Supporting information

Supplemental script 1

Supplemental script 2

## ACKNOWLEDGEMENTS

We thank the LIC Imaging Center Freiburg for expertise and support. *This project is funded by the Deutsche Forschungsgemeinschaft (DFG, German Research Foundation):* CIBSS – EXC2189 Project ID 390939984 (to E.B.) and confocal microscopy *Project IDs* 414136422.

## SUPPLEMETARY INFORMATION

*File: Supplementary scripts_Roessling_Absolute apoplastic pH:*

Supplementary Script 1: HPTS Ratiometry Script (Updated from Barbez et al., 2017) Supplementary Script 2: Ratio to Absolute pH Conversion (Version 20250606)

