## Supplemental script 2 for "A workflow for absolute apoplastic pH assessment during live cell imaging in plant roots"

Supplemental script 2: Ratio to Absolute pH Conversion: 
Developed in this work and available Zenodo: https://doi.org/10.5281/zenodo.15599806

This Fiji Jython script converts 465/405 ratiometric images of HPTS-stained Arabidopsis roots into absolute apoplastic pH maps. It uses a min�max normalization approach and a polynomial calibration function derived from in vitro HPTS data. The script enables high-resolution, pixel-wise quantification of extracellular pH. We recommend downloading the script from Zenodo where always the newest plugin version will be available. 
Features:
* Converts 32-bit grayscale ratio images into absolute pH values.
* Supports two calibration modes:
� Manual: enter lower and upper ratio values (e.g., from pH 5.0 and 7.0 samples)
� From images: use calibration images to automatically determine normalization values
* Applies a user-defined polynomial equation (up to third order) for pH conversion.
* Outputs 32-bit TIFF images with pH-based LUT visualization.
Usage:
1. Open Fiji, create a new script (Plugins > New > Script...), set the language to Jython.
2. Copy and paste the script code into the editor.
3. Run the script and fill in the input parameters 
4. Ensure that calibration and sample images were acquired with identical confocal settings.
5. Resulting images will be saved to the specified output folder, with pixel values representing absolute pH.

For questions or feedback, contact:
Elke Barbez, University of Freiburg,


# -------------------------------------------------------------
# Script Name: Ratio to absolute pH conversion
# Version: 5
# Description:
#     Converts 465/405 ratiometric images to absolute apoplastic pH using
#     user-defined normalization and polynomial conversion.
#     Users must choose one calibration method: manual or from image means.
#     Saves output as 32-bit TIFFs with LUT visualization.
# -------------------------------------------------------------

#@ String(value="<html style=\"width: 400px;text-align: center;\"><p style=\"margin:0px;padding:0px;font-size:12px;\"><b>Ratio to Absolute pH Conversion</b></p><p>This script converts ratiometric images (465/405) into absolute apoplastic pH values using a polynomial calibration. Choose either manual input or calibration images.</p></html>", visibility=MESSAGE, required=false) desc
#@ File(label="Folder with your images (?)", style="directory", description="Input folder") input_dir
#@ File(label="Folder to save your images (?)", style="directory", description="Output folder") output_dir
#@ String(label="Extension for the images to look for (?)", value="tif", description="Extension of your images to select in the input folder") extension_ratio
#@ String(value="<html style=\"width: 233px;\"><div style=\"height: 1px; background: #c0c0c0;\"/></html>", visibility=MESSAGE, required=false) line1
#@ String(label="Calibration mode", choices={"Manual", "From calibration images"}, value="Manual") calib_mode
#@ Double(label="Lower calibration ratio (lower pH)", stepSize=0.000001, required=false) lower_ratio
#@ Double(label="Upper calibration ratio (upper pH)", stepSize=0.000001, required=false) upper_ratio
#@ File(label="Lower calibration image (lower pH)", style="file", required=false) lower_image
#@ File(label="Upper calibration image (upper pH)", style="file", required=false) upper_image
#@ String(value="<html style=\"width: 233px;\"><div style=\"height: 1px; background: #c0c0c0;\"/></html>", visibility=MESSAGE, required=false) line2
#@ Double(label="B3 (coefficient for x? )", stepSize=0.0001, value=3.4347) B3
#@ Double(label="B2 (coefficient for x? )", stepSize=0.0001, value=-5.7843) B2
#@ Double(label="B1 (coefficient for x? )", stepSize=0.0001, value=4.2768) B1
#@ Double(label="B0 (constant term)", stepSize=0.0001, value=5.0497) B0
#@ String(value="<html style=\"width: 233px;\"><div style=\"height: 1px; background: #c0c0c0;\"/></html>", visibility=MESSAGE, required=false) line3
#@ String (label="LUT for visualization (?)", choices={"Green Fire Blue", "Fire", "Grays", "Ice", "Spectrum", "Red", "Green", "Blue", "Cyan", "Magenta", "Yellow", "Red/Green", "Cyan Hot", "HiLo", "ICA", "ICA2", "ICA3", "Magenta Hot", "Orange Hot", "Rainbow RGB", "Red Hot", "Thermal", "Yellow Hot", "blue orange icb", "cool", "gem", "glow", "mpl-inferno", "mpl-magma", "mpl-plasma", "mpl-viridis", "phase", "physics", "royal", "sepia", "smart", "thal", "thallium", "unionjack"}, description="Select the lookup table for final coloring") lut_method
#@ Double(label="pH range minimum", value=5.0, stepSize=0.01) pH_min
#@ Double(label="pH range maximum", value=7.0, stepSize=0.01) pH_max
#@ String(value="<html style=\"width: 400px;text-align: center;\">Please cite Barbez et al. 2017<br/>and R��ling et al. 2025</html>", visibility=MESSAGE, required=false) footer

# ??? IMPORTS ????????????????????????????????????????????????????????????????????

import os
import math
import time
import fnmatch

from ij import IJ, ImagePlus
from ij.process import FloatProcessor
from ij.plugin import LutLoader

# ??? FUNCTIONS ??????????????????????????????????????????????????????????????????

def progress_bar(progress, total, line_number, prefix=""):
    """Progress bar for the IJ log window

    Parameters
    ----------
    progress : int
        Current step of the loop
    total : int
        Total number of steps for the loop
    line_number : int
        Number of the line to be updated
    prefix : str, optional
        Text to use before the progress bar, by default ''
    """

    size = 30
    x = int(size * progress / total)
  
    IJ.log(
        "\\Update%i:%s\t[%s%s] %i/%i\r"
        % (line_number, prefix, "#" * x, "." * (size - x), progress, total)
    )


def timed_log(message):
    """Print a log message with a timestamp added

    Parameters
    ----------
    message : str
        Message to print
    """
    IJ.log(time.strftime("%H:%M:%S", time.localtime()) + ": " + message)


def getFileList(directory, extensions):
    """Get a list of files with the extension

    Parameters
    ----------
    directory : str
        Path of the files to look at
    extensions : [str]
        Extensions to look for

    Returns
    -------
    list
        List of files with the extension in the folder
    """
    
    files = []
    filteringStrings = [ext.lower() for ext in extensions]
    for dirpath, dirnames, filenames in os.walk(directory):
        for filename in filenames:
            # Check if the file matches any of the extensions (case-insensitive)
            if any(fnmatch.fnmatch(filename.lower(), "*" + ext) for ext in extensions):
                files.append(os.path.join(dirpath, filename))
    return files


def normalize_pixel(val, lower, upper):
    """Normalizes a pixel value within a given range

    Parameters
    ----------
    val : float
        The pixel value to be normalized
    lower : float
        The minimum value of the normalization range
    upper : float
        The maximum value of the normalization range
        
    Returns
    -------
    float
        The normalized value 
    """

    norm = (val - lower) / (upper - lower)
    return max(0.0, min(1.0, norm))  # Clamp to [0, 1]


def convert_to_pH(norm_val, B3, B2, B1, B0):
	"""Converts a normalized value to a pH value using a polynomial

    Parameters
    ----------
    norm_val : float
        The normalized value (between 0.0 and 1.0) that will be transformed into a pH value
    B3 : float
        The coefficient for the cubic term (x?) 
    B2 : float
        The coefficient for the quadratic term (x?) 
    B1 : float
        The coefficient for the linear term (x?) 
    B0 : float
        The constant term (the offset) 
        
    Returns
    -------
    float
        The calculated pH value based on the polynomial: B3 * x? + B2 * x? + B1 * x + B0.
    """
    
	return B3 * norm_val**3 + B2 * norm_val**2 + B1 * norm_val + B0


def process_image(img, lower, upper, B3, B2, B1, B0):
	"""Processes an image to convert pixel values into pH values by normalizing and applying a polynomial.
	
	Parameters
	----------
	img : ImagePlus
	    The input image whose pixel values will be processed and converted into pH values
	lower : float
	    The lower bound for the normalization of pixel values
	upper : float
	    The upper bound for the normalization of pixel values
	B3 : float
	    The coefficient for the term (x?) used in the conversion to pH
	B2 : float
	    The coefficient for the term (x?) used in the conversion to pH
	B1 : float
	    The coefficient for the term (x?) used in the conversion to pH
	B0 : float
	    The constant term (offset) used in the conversion to pH.
	
	Returns
	-------
	ImagePlus
	    A new imageplus where pixel values are converted into pH values
	"""
    
   	ip = img.getProcessor().duplicate().convertToFloat()
	width = ip.getWidth()
   	height = ip.getHeight()
   	pH_ip = FloatProcessor(width, height)
   	
   	for y in range(height):
   		for x in range(width):
   			val = ip.getf(x, y)
   			
   			if math.isnan(val) or val == 0.0:
   				pH_ip.setf(x, y, float('nan'))
   			else:
   				norm_val = normalize_pixel(val, lower, upper)
   				pH_val = convert_to_pH(norm_val, B3, B2, B1, B0)
   				pH_ip.setf(x, y, pH_val)

   	return ImagePlus("pH_" + img.getTitle(), pH_ip)


def get_mean_intensity(image_path):
    """Gets the mean intensitiy of an ImagePlus 
	
	Parameters
	----------
	image_path : String
	    File path of the calibration image to be used

	Returns
	-------
	float
	    The mean value 
	"""
    
    img = IJ.openImage(image_path)
    if img is None:
        timed_log("Error: Cannot open calibration image: {}".format(image_path))
        raise ValueError("Cannot open calibration image: {}".format(image_path))
    if img.getProcessor().getBitDepth() != 32:
        timed_log("Error: Calibration image is not 32-bit: {}".format(image_path))
    	raise ValueError("Calibration image is not 32-bit: {}".format(image_path))
    	    
    stats = img.getStatistics()
    return stats.mean
    
# ??? MAIN CODE ??????????????????????????????????????????????????????????????????

IJ.log("\\Clear")
timed_log("Script starting")


if calib_mode == "From calibration images":
    if lower_image is None or upper_image is None:
        timed_log("Error: Both calibration images must be provided.")
        raise Exception("Both calibration images (lower pH and upper pH) must be selected for image-based calibration.")
    lower_ratio = get_mean_intensity(lower_image.getAbsolutePath())
    upper_ratio = get_mean_intensity(upper_image.getAbsolutePath())
    IJ.log("") # Place for progress bar
    timed_log("Info: Calibration image (lower pH) -> mean ratio: {:.6f}".format(lower_ratio))
    timed_log("Info: Calibration image (upper pH) -> mean ratio: {:.6f}".format(upper_ratio))

elif calib_mode == "Manual":
    if lower_ratio is None or upper_ratio is None:
        timed_log("Error: Both manual calibration values must be entered.")
        raise Exception("Manual calibration mode requires both lower and upper values.")

else:
    timed_log("Invalid calibration mode selected.")
    raise Exception("Invalid calibration mode.")


input_dir = input_dir.getAbsolutePath()
output_dir = output_dir.getAbsolutePath()

file_ext_filter = [extension_ratio]
files = getFileList(input_dir, file_ext_filter)
total_files = len(files)


for i, file in enumerate(files, 1):
    # Get basename of file
    basename = os.path.basename(file) 
    
    progress_bar(i, total_files, 1, "Processing: " + str(i))

    img = IJ.openImage(file)    
    if img is None:
        timed_log("Info: Could not open image: {}".format(basename))
        continue
    
    if img.getProcessor().getBitDepth() != 32:
    	timed_log("Info: Image is not 32-bit: {}".format(basename))
    	continue

	# Process the image 
    pH_img = process_image(img, lower_ratio, upper_ratio, B3, B2, B1, B0)
    
    # Apply LUT
    if not LutLoader.getLut(lut_method):
    	# Select LUT does not exist. 
    	# Check if Green Fire Blue exists 
        if LutLoader.getLut("Green Fire Blue"):
            timed_log("LUT '" + lut_method + "' does not exist. Using default: 'Green Fire Blue'")
            lut_method = "Green Fire Blue"
        else:
        	# Green Fire Blue does not exist. 
        	# Use a built-in LUT
            timed_log("LUT '" + lut_method + "' does not exist. Using default: 'Fire'")
            lut_method = "Fire"
    IJ.run(pH_img, lut_method, "")
    
    # Set Calibration Bar
    IJ.setMinAndMax(pH_img, pH_min, pH_max)
    IJ.run(pH_img, "Calibration Bar...", "location=[Upper Right] fill=White label=Black number=5 decimal=3 font=12 zoom=1 overlay")

	# Save image
    out_path = os.path.join(output_dir, pH_img.getTitle())
    IJ.saveAs(pH_img, "Tiff", out_path)

timed_log("Script finished !")
print("Script finished !")
